# Evolutionary Genetic Species Detected in Prokaryotes by Applying the K/θ Ratio to DNA Sequences

**DOI:** 10.1101/2020.04.27.062828

**Authors:** C. William Birky, Heather Maughan

**Affiliations:** Department of Ecology and Evolutionary Biology, The University of Arizona, Tucson, Arizona 85721, United States of America; Ronin Institute, Montclair, New Jersey 07043, United States of America

**Keywords:** bacterial species, evolutionary genetic species concept, K/θ species criterion

## Abstract

Previous papers in this series described the “evolutionary genetic species concept” which is based on population genetic theory and should be applicable to any organism. Also described was a species criterion, the K/θ ratio, that delimits independently evolving evolutionary species based on single-gene sequences, and its application to sexual and asexual eukaryotes was illustrated. Here, we show how the evolutionary genetic species concept and the K/θ ratio can be applied to bacteria, using the sequences from some genes of the core genome that are rarely, if ever, transferred horizontally between different species. This completes the demonstration that K/θ is a general method for species delimitation, applicable to all organisms. Also, it adds to the evidence that bacteria have species in the most general sense, even though they have the ability to exchange genes across species boundaries. Finally, we show that a published critique of the use of K/θ ≥ 4 as a criterion for independently evolving species rests on two errors in the application of population/evolutionary genetic theory.

## Introduction

A prominent feature of the biological world is discontinuous variation, with clusters of similar organisms differing significantly from organisms in other clusters. These clusters, called species and given different binomial names, were originally recognized by their different phenotypes. With the development of population genetic theory, species have been recognized as populations with distinct gene pools underlying their phenotypic differences. When gene sequences are used to reconstruct evolutionary history in the form of phylogenetic trees, different species appear as different clades of similar sequences separated by deep gaps in the tree.

This background is widely, but not universally, accepted. Doubters often focus on the gradual divergence of sister populations from each other, wondering at what point they should be called different species. Even among those who agree that species are real, distinct populations, there is disagreement on how to define and recognize them, given the gradual divergence of recently separated populations. Many authors simply point to “deep gaps” in a phylogenetic tree, usually of gene sequences, as evidence for different species, but this raises the question: “How deep is deep enough?”

This question can be answered by thinking of species as independently evolving populations. This is arguably the most general model of species (species concept), applicable in principle to all organisms and consistent with population and evolutionary genetic theory [1-8]. Different species are populations that evolve independently of each other, in the sense that they are independent arenas for mutation, natural selection, and random genetic drift. In other words, a new mutant allele arising in one population cannot spread to the other, or if it does move from one species to the other by hybridization, it is very unlikely to be fixed in that population. Moreover, if a shared allele is fixed or lost in one population by drift and/or selection, this does not affect the probability that it will be fixed or lost in the other. This is referred to as the Evolutionary Genetic Species Concept (EGSC) to distinguish it from the Evolutionary Species Concept (EvSC) of Simpson [7]. The EvSC as it is usually worded requires that we know, or assume, that the populations in question will continue to evolve independently in the indefinite future. The requirement for future independence should not be part of a species concept because it is unknowable; for instance, a trait difference required for adaptation of species A to a different niche from species B could be lost in the future. (However, it should be noted that in 1953 Simpson did allow that speciation might occur in allopatry without ecological differences in “some probably rare cases” [[7], p. 171]).

In practice, a pair of sister species are recognized as soon as they have diverged to the point where samples of individuals can be shown to come from two independently evolving populations. This can be done using a number of different criteria for independent evolution. Prominent examples that have been applied to prokaryotes as well as eukaryotes are reproductive isolation and ecological isolation, both of which obviously result in independent evolution of the isolated populations. However, both of these species criteria have conceptual and practical problems. Neither one is generally applicable; reproductive isolation is relevant only for organisms with significant amounts of sexual reproduction, and neither criterion includes the possibility that species may be evolving independently simply because they are physically isolated from each other. Moreover, neither is readily applicable to large-scale biodiversity surveys or species discovery that relies on only sequences, as in the Barcode of Life [9] or Multi-Locus Sequence Typing projects [10].

Although DNA sequences are widely used to delimit species in prokaryotes, there is a great deal of controversy about how this should be done, and even about whether bacteria really have species. Some of this controversy mirrors similar debates about species in eukaryotes. However, much of it arises from the phenomenon of horizontal gene transfer (HGT), in which segments of genes, whole genes, and even large segments of chromosomes are transferred between distantly related bacteria via common mechanisms of bacterial gene exchange: conjugation, transformation, and transduction. This problem can be solved by distinguishing between core genes and auxiliary genes, as core genes are less likely to be transferred or retained if transferred [11, 12]. Furthermore, HGT doesn’t appear to affect the bacterial species identification method based on average nucleotide identity (ANI), where the nucleotide identity of each pair of orthologous genes from two genomes is calculated and then averaged over all genes shared between the two genomes [13]. The use of numerous genes is expected to diminish the effects of HGT on sequence-based approaches to delimiting species. Although ANI-based methods have proven to be useful, they are not based on theory but derive from empirical observations.

In this paper, as in previous publications [1-6], we argue that the EGSC can be applied to all organisms, sexual and asexual, prokaryote and eukaryote. We also show how the K/θ ratio, first used to discover and delimit EG species in eukaryotes based on DNA sequence data from a single gene, can be applied to bacteria if a core protein-coding gene or 16S rRNA gene is used. In particular, it could be applied to large-scale diversity surveys of bacteria using such a gene. Such surveys are routine for eukaryotes, and can now be done with bacteria regardless of whether they can be cultured in the lab, thanks to the development of methods that allow amplification of genes from single bacterial cells isolated by dilution or flow cytometry from environmental samples (e.g. [14, 15]]. The successful identification of evolutionary genetic species in bacteria using the K/θ ratio (this paper) as well as methods such as the Generalized Mixed Yule Coalescent (GMYC; [1, 16]) adds to the growing literature that demonstrates that EG species are a general phenomenon found in varying degrees in all organisms, sexual and asexual, prokaryotes and eukaryotes.

The K/θ test is designed to distinguish between two kinds of clades: (i) those separated by shallow gaps formed within a single species by random genetic drift or transient physical isolation, and (ii) those separated by deep gaps produced by adaptation to different ecological niches and/or by long-term physical isolation, i.e. different species. Simulations [17] and population genetic theory [18] show that 95% of the gaps between clades within a species are expected to have a depth of < 4N_e_ generations (where N_e_ is the effective population size), while gaps between species clades can be of any depth. It is thus possible to recognize a pair of clades as different species, with probability ≥ 0.95, if they are separated by gaps of ≥ 4N_e_ generations. At that point the sequence difference between the two clades is *K* ≥ 8*N*_*e*_*μ* (where μ is the mutation rate per base pair) and the ratio K/θ ≥ 8N_e_μ/2N_e_μ ≥ 4 (3-8). (Barraclough’s [19] claim to have found errors in this theory is erroneous, as we show below in Discussion.)

This result is based on expected probabilities and does not account for uncertainty introduced by finite sample sizes. The relevant sampling theory has been developed by Noah Rosenberg [20, 21], who showed that it holds for any pair of sister clades, i.e. reciprocally monophyletic samples n1 and n2, provided the sample sizes are at least n1 = 3 and n2 = 2. The power of these very small samples may seem surprising, but it arises from their first being shown to be reciprocally monophyletic.

To illustrate how the K/θ ratio is used to detect evolutionary genetic species in prokaryotes, we use the set of sequences of six different housekeeping/core genes previously used by Wertz et al. [22, 23] to delimit species in enteric bacteria. This was one of the first multilocus data sets used to discover bacterial species. It provides an opportunity to apply the K/θ method to single genes and compare the results of using different genes to each other and to the named taxonomic species defined by classical phenotypic traits. Note that these data are used simply to demonstrate the application of the K/θ method and are not intended to be definitive statements about these specific bacterial species.

## Materials and Methods

John Wertz kindly provided partial sequences of six housekeeping core genes (*gapA, groEL, gyrA, ompA, pgi*, and 16S rRNA) from seven named species of enteric bacteria (*Citrobacter freundii* (abbreviated CF), *Enterobacter cloacae* (EB), *Escherichia coli* (EC), *Hafnia alvei* (HA), *Klebsiella oxytoca* (KO), *Klebsiella pneumonia* (KP), *Serratia plymuthica* (SP), and the outgroup *Vibrio cholera* (VC)).

The aligned sequences for each gene were visualized in MacClade [24] where all the sequences of the gene were trimmed to the same length and all internal sites with gaps in any one sequence were removed. Consequently, all pairwise sequence differences for a given gene are based on the same regions of the gene. Final sequence lengths were: *gapA*, 540 bp; *groEL*, 348 bp; *gyrA*, 634 bp; *ompA*, 433 bp; *pgi*, 242 bp; *16S*, 289 bp. One internal gap of 27 bp in the *ompA* gene of the outgroup *V. cholerae* was retained because to remove it from all sequences would cause an unnecessary loss of sequence information in the ingroup.

To discover species using the trimmed sequences, we followed the procedure outlined by Birky [2] for each gene separately. Briefly,

1. First, we used PAUP* [25] to find a bootstrapped Neighbor-joining tree in which all clades were supported by ≥70% of 1000 bootstrap replicas. A Neighbor-joining tree was used to rearrange the sequences in the PAUP* file to reflect the order of sequences in the tree; this greatly simplifies subsequent calculations in Excel, but note that the tree itself is not used in subsequent calculations. Both corrected and uncorrected sequence differences were then exported from PAUP* to an Excel spreadsheet where all subsequent calculations were performed. Starting at the tips of the tree, each pair of sister clades is tested separately in steps 2-7, to determine the probability that the clades are samples from independently evolving species. This process is continued with deeper and deeper clades until species are found. For clades that are members of a non-bifurcating tree structure such as a polytomy (A, B, C) or ladder ((A,B)C), we compare A and B first, then compare C to whichever one of those is closest to C.
2. For the two candidate clades in a pair, the mean pairwise difference *d* between sequences is multiplied by the sample size correction n/(n-1) where *n* is the number of sequences in the clade, to get the nucleotide diversity *π*. Note that nucleotide diversity is defined as a function of the uncorrected pairwise distance. When *d* = 0, we used a non-zero estimate of *π* by assuming that one sequence differs from the others by 1/L where L is the sequence length; then *π* = 2/Ln(n-1).
3. For each clade, we then estimate θ = 2N_e_μ by *π* /(1 -4 *π* /3).
4. Calculate K = mean pairwise sequence difference between the two clades, corrected for multiple substitutions.
5. Calculate K/θ for the pair of clades. When the values of θ for the two clades differ, we use the larger value to get a conservative estimate of K/θ. If K/θ ≥ 4, we conclude that the individuals in those clades came from different species, with probability ≥ 0.95.
6. The preceding steps are based on expected probabilities and do not account for uncertainty introduced by finite sample sizes. This uncertainty is considered as follows, using results obtained by Noah Rosenberg [20]. Using the K/θ ratio and the numbers *n*1 and *n*2 of individuals in the two clades, we want to find the probability that the individuals were sampled from populations that have been evolving independently long enough to become reciprocally monophyletic. This is best done using a table available on request from Noah Rosenberg or the first author; alternatively, the values can be estimated from Figure 6 in Rosenberg [20].

It is worth repeating that the phylogenetic tree used in step 1 is only used for two purposes: (1) to facilitate the identification of sister clades, and (2) to inform the organization of the Excel spreadsheet of pairwise sequence differences with the most similar sequences together. The calculations could be done without the tree, which would require identifying pairs of most similar sequences by eye in the spreadsheet. Thus, the K/θ method is not affected by discrepancies between distance-based phylogenetic inference methods.

## Results

The Wertz et al. [23] data consist of 5 protein-coding genes plus part of the 16S rRNA gene. We will first illustrate the application of the K/θ ratio by showing the detailed analysis for *pgi*, the protein-coding gene that resolves the most species. The high resolution suggests that this gene segregated before most of the other genes in most of the speciation events, by chance. We will then summarize the results obtained with each of the other genes. Following Wertz et al. [23], we consider the single isolate of *S. marcescens* as belonging to the same taxonomic species as *S. plymuthica* because for all the genes in this study it is identical or nearly so to the *S. plymuthica* isolates.

### Finding species with *pgi*

The Neighbor-joining (NJ hereafter) tree in Figure 1 shows that the uncorrected *pgi* sequences form a number of clades that are seen in ≥ 63% of 1000 bootstraps; most have 100% support. The taxonomic species *K. oxytoca*, KO; *E. cloacae*, EB; *H. alvea*, HA; and *S. plymuthica* SP, SM all form clades with 100% bootstrap support, but some of these in turn contain two or more strongly supported clades. We applied the K/θ test to each well-supported pair of sister clades, starting at the bottom of the *pgi* tree in Figure 1. These clades are also well-supported in the NJ tree using corrected sequence differences. The Excel table for *H. alvea* and *S. plymuthica* is shown in Table 1 as an example of the data. The full tables for all genes are available on request from the first author.

**Table 1.** Matrix of pairwise sequence differences for *pgi* from *Serratia plymuthica* and *Hafnia alvea*.

|  | SM1 | SP1 | SP2 | HA1 | HA2 | HA3 | HA4 |
| --- | --- | --- | --- | --- | --- | --- | --- |
| SM1 | - |  |  |  |  |  |  |
| SP1 | 0.0000 | - |  |  |  |  |  |
| SP2 | 0.0000 | 0.0000 | - |  |  |  |  |
| HA1 | **0.3022** | **0.3022** | **0.3022** | - |  |  |  |
| HA2 | **0.3010** | **0.3010** | **0.3010** | 0.0083 | - |  |  |
| HA3 | **0.3087** | **0.3087** | **0.3087** | 0.0125 | 0.0042 | - |  |
| HA4 | **0.3010** | **0.3010** | **0.3010** | 0.0083 | 0.0000 | 0.0042 | - |

Uncorrected sequence differences $d$ and corrected differences $K$ (bold type).

**Figure 1.**
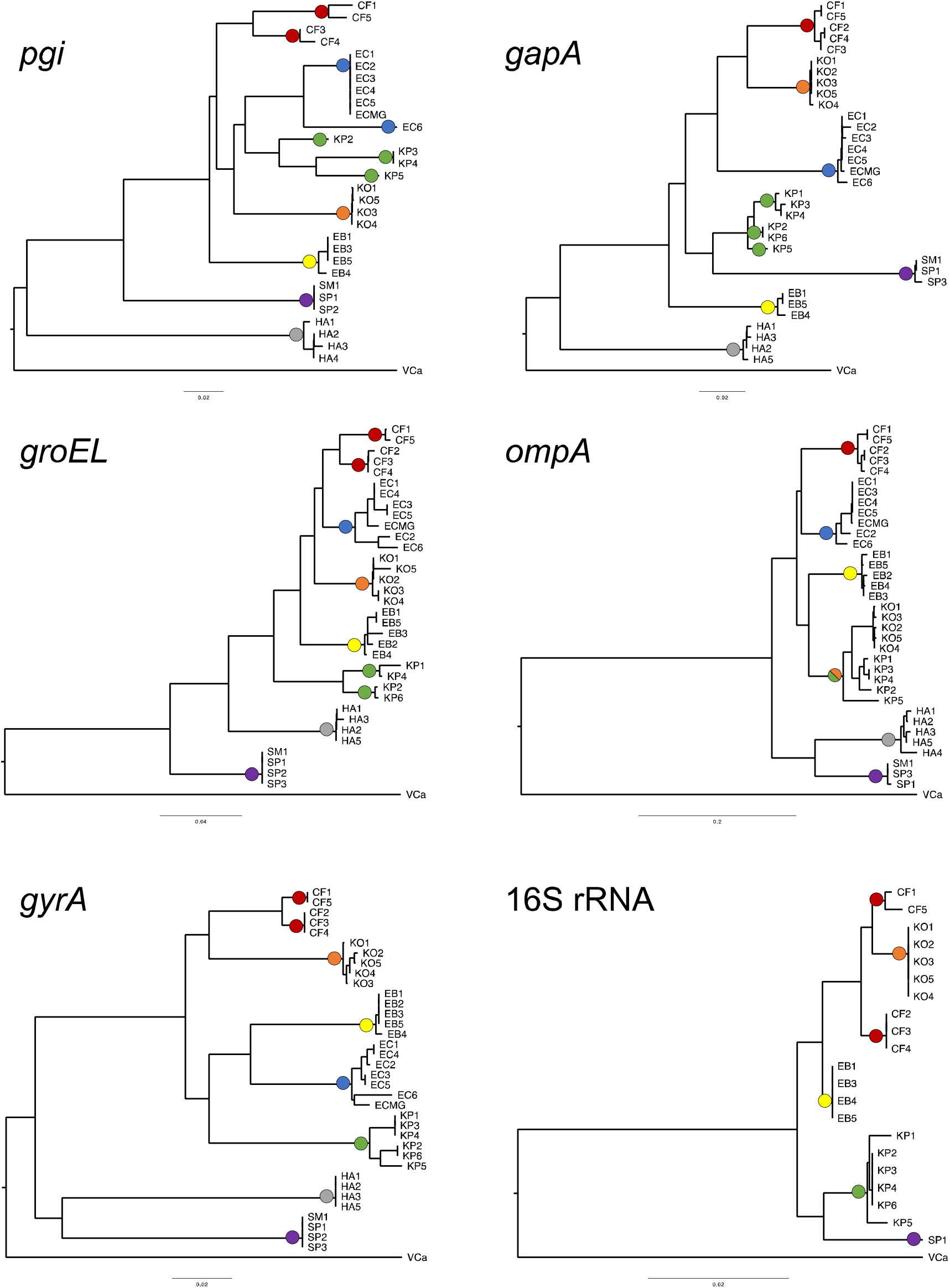
NJ trees of *pgi, groEL, gyrA, gapA, ompA* or 16S rRNA sequences. CF, EC, KP, KO, EB, SM, and HA are taxonomic species. Closed circles indicate clades that are different EG species delimited by the K/θ method, with circle colors representing different taxonomic species. All scale bars indicate 0.02 substitutions per site except for those of *groEL* and *ompA*, which represent 0.04 and 0.2 substitutions per site, respectively.

1. To illustrate the procedure, we begin with clade HA2,4, which has weak support (69% of bootstraps); its sister group is the singlet HA3. Using the formula for a maximum estimate of *π* and θ when d = 0, we calculate that θ = 0.00836 and from the table K = 0.0021 so K/θ = 0.25 and these are not different species. This result was obvious from inspection and normally would not warrant a calculation. Next we ask if clade HA1-4 is a different species from clade (SM1,SP1,SP2). The pairwise distances for SM1, SP1, and SP2 are zero while those for HA1-4 are not all zero; we prefer to use the latter because they do not require any correction. For this clade d = 0.00625, *π* = 0.00833, and θ = 0.00843. K is 0.03032 so K/θ = 36.0 and these are samples from different clades with probability > 0.988. Because K/θ >> 4, these specimens are also samples from different independently species with P >> 0.95. Note that these results are obvious by inspection of the branch lengths in an NJ tree like that shown for *pgi* in Figure 1 in which branch lengths are proportional to the number of substitutions. The branches connecting the (SM,SP) clade to the HA clade are clearly more than four times the length of the branches connecting any two specimens within the HA clade.
2. The taxonomic species *C. freundii* (CF) forms two clades, CF1,5 and CF3,4, each with 100% bootstrap support. Using Excel, we calculate the mean corrected sequence difference between these clades, K = 0.07618. CF1,5 has the larger value of θ = 0.0437, giving K/θ = 1.74. In Rosenberg’s table, when K/θ = 1.7 and the sample sizes are 2 and 2, the probability that these two clades are samples from reciprocally monophyletic populations is only 0.57. Moreover, because K/θ is much less than 4, the probability that the samples came from different species is low. This is the more conservative test in terms of splitting species because it uses the larger of the two values of θ. On the other hand, the clade CF3,4 has the smaller θ = 0.00167, giving K/θ = 4.5. From Rosenberg’s table, *P* = 0.97 that these are samples from different clades; furthermore, because K/θ > 4, the probability is very high that they are from different independently evolving species. However, this is based on the less conservative estimate of θ. Because only one of the two possible estimates of θ support the hypothesis of different species, we conclude that the *pgi* gene shows weak support for two cryptic species within *C. freundii*. As we will see below, some, but not all, other genes strongly support splitting. This suggests that there is enough recombination in *C. freundii* to cause different genes to complete segregation at different times, by chance.
3. The taxonomic species *E. coli* (EC) also consists of a clade (EC1-5,ECMG) with 100% bootstrap support and a singleton EC6. Only the clade can be used to estimate θ, but since the 6 sequences in the clade are identical, π and thus θ cannot be calculated directly. For these cases, one can conservatively estimate these values by assuming that one of the 6 sequences actually differs from the others at a single site. Then using the calculation given in Materials and Methods, one can calculate π = 0.001377, θ = 0.00166, and K = 0.07000 so K/θ = 42.3 and there is strong evidence that the clades EC1-5,MG and EC6 are samples from different clades and from different cryptic species. This procedure is likely to overestimate π and θ and hence underestimate K/θ, so in this sense it is conservative with respect to splitting species.
4. Next we consider the clade consisting of *pgi* from KP isolates. Starting at the tips, we compared (KP3,4) with KP5. Using the same procedure as above to get a non-zero value for π, K/θ = 8.38, providing evidence that these are samples from two different clades and species. Comparing (KP3,4) to the other singleton, KP2, we find K/θ = 11.6 and these are also from different species. Thus KP is divided into three species by these gene sequences.
5. Finally, the sequence data from this gene confirm that the isolates identified as *K. oxytoca* (KO) and *E. cloacae* (EB) are evolutionary genetic species.

To summarize, the sequences of *pgi* confirm that KO, HA, EB, and SM,P are species; split EC into 2 species; and split KP into 3 species.

### Finding species with *groEL*

The *groEL* gene (Figure 1) confirms that KO, EB, SM, and HA are species. It splits CF into two species, CF1,5 and CF3,4. EC is confirmed as a separate species from its sister clade CF2-4 if one uses θ from CF2-4, but not if one uses θ from EC. *groEL* sequences split KP into KP1,4 and KP2,6 if one uses θ from KP2,6; however, they do not make this split if one uses the larger θ from KP1,4.

### Finding species with *gyrA*

The gene *gyrA* (Figure 1) confirms that KO, EB, EC, HA, and SM are Evolutionary Genetic Species. The sequences of this gene split the taxonomic species CF into two EG species, CF1,5 and CF2-4, and also split KP into KP1,4 and KP2,6.

### Finding species with *gapA*

The *gapA* gene (Figure 1) confirms that CF, KO, EC, SM,SP, EB, and HA are different species. KP consists of two clades, KP1,3,4 and KP2,6, and a singlet KP5 which are samples from three different species.

### Finding species with *ompA*

The *ompA* gene (Figure 1) provides evidence that the CF, EC, EB, HA, and SP,SM sequences each represent a single species. Unlike some of the other genes, it does not split any of these species. In contrast, the KO and KP sequences are polyphyletic. Although the KO sequences are very similar with a diversity commensurate with that of other species in these data, the KP sequences are very divergent from each other. Moreover, KP5 is basal to KP1-4 as well as to KO; thus KP is polyphyletic, suggestive of incomplete lineage sorting.

### Finding species with 16S

The 16S sequences of the taxonomic species *E. coli* and *H. alvia* contained regions that were not present in the other species and for which we could find no explanation, so these two taxonomic species were omitted from the analysis of the 16S sequences. Also, all pairwise differences among the ingroup sequences were ≤ 0.063, including those between species, so no corrections for multiple hits were applied. The K/θ test verified that the CF1,5, KO, CF2-4, EB, and KP sequence sets, and the SP1 singlet, each represent evolutionary genetic species with probabilities >0.99. However, the CFl and CF5 sequences did not; the clade was supported by only 56% of bootstrap replicas, so no K/θ test was done. The NJ tree is shown in Figure 1.

### Comparison with ANI

To compare these results with ANI values we used *d*, the mean pairwise difference for all bacterial sequences for each gene. ANI is equal to 1-*d*, multiplied by 100 to make it a percentage. The resulting values for the genes included in our analysis ranged from 95.9 to 100% ANI, all greater than the 95% cutoff for within-species differences [26]. Note that these are values for single genes, not whole genomes, but if each gene is representative of its genome, this means that species delimited with the K/θ ≥ 4 criterion will also be identified as species by the accepted ANI criterion.

## Discussion

Our focus here, and in several preceding papers on discovering species in asexual and sexual eukaryotes [1-6], is on two problems: (i) identifying a model of species, i.e. a species concept, that is general enough to cover all organisms, whether prokaryotes or eukaryotes, and all degrees and kinds of sexuality; and (ii) based on the model, testing a general method of assigning specimens to the same or different species (a species criterion), using gene sequences. We showed that the Evolutionary Genetic species concept and the K/theta ratio can be used to delimit bacterial species using DNA sequences, and that this procedure often splits species defined by traditional systematics.

The results with the protein-coding genes and isolates EB, EC, HA, KO, and SM/P are straightforward: each of the genes shows that these five taxonomic species, discovered by traditional methods of bacterial taxonomy using morphology and various phenotypic tests, are also EG species. The situation with CF is more complex. All five genes show two clades, (CF1,5) and (CF2-4) with 89-100% bootstrap support except *gapA* where the support for the (CF2-4) clade is only 64%. Moreover the K/θ ratio for this pair of clades is greater than 4, signifying different species, for *gyrA, pgi*, and *groEL* but not for *gapA* or *ompA*.

### Deep gaps in trees separating species: how deep is deep enough?

In the literature of new species descriptions, and in papers proposing the splitting of species, it is common to refer to deep gaps between clades in phylogenetic trees as evidence that the clades should be assigned to different species. The decision as to whether the gap between two clades A and B is deep enough to assign A and B to different species, as opposed to indicating a relatively shallow gap within a single species, is generally made using the systematist’s intuition. Occasionally the systematist compares the gap between A and B to the gap between named species C and D, concluding that A and B are different species if the gap between them appears to be as deep or deeper than the gap between C and D. However, this assumes that species C and D are well defined. The K/θ method avoids making these assumptions by providing a decision criterion that is soundly based on population genetic theory.

### Barcode gaps

If one plots a frequency distribution of pairwise sequence differences between the DNA sequences of a set of specimens, one often finds a bimodal distribution; the space between the modes is called a barcode gap. This is the basis of the Barcode of Life project. Eckert and colleagues [27] used nearly complete 16SrRNA sequences to look for a barcode gap in Cyanobacteria, using the Automatic Barcode Gap Discovery (ABGD) program; they found gaps in some but not all taxonomic groups. However, rather than testing pairs of sister clades, Eckert et al. pooled between 34 and 2448 sequences from each of 16 genera. This will obscure the barcode gap if different pairs of species have gaps in different places, i.e. if some species have values of θ that overlap the values of K. It is also possible that some of the groups examined by Eckert et al. have undergone a recent adaptive radiation such that K values are very small, as the authors noted. In contrast, Jain et al. [28] found a pronounced gap in plots of ANI from 91,761 whole genome sequences from GenBank; the gap stretched from 83-95% ANI and contained only 0.2%. of the pairwise comparisons.

### Not all well-supported clades are species

These analyses illustrate the danger of assuming that all clades are species. For example, in the *pgi* tree shown in Figure 1 the clade EB1,3,5 has 90% bootstrap support and yet belongs to the same species as EB4. If one studied *E. cloacae* EB only, one could easily draw a tree with the Y axis stretched out, so that the branch connecting the clade to EB4 looks very long and might appear to show two species connected by a long branch.

### Applicability of the K/θ method

It is clear that the Evolutionary Genetic Species Concept is applicable to bacteria. It is also clear that within the framework of the EGSC, species can be discovered using DNA sequences analyzed with the K/θ method. It is important to note that the K/θ test was deliberately designed to work with a single gene or sequence. If the sequences of two or more genes are used, it is important to remember that in a sexual organism (including bacteria with high levels of recombination) different genes may become reciprocally monophyletic at different times. We recommend that the K/θ test be applied to each gene separately; if any gene places the specimens in two different clades and K/θ ≥ 4, this would be strong evidence that the clades represent different species. We do not recommend concatenating gene sequences without first testing whether the signal from a gene that has not yet segregated can obscure the signal from a gene that has become reciprocally monophyletic, i.e. could cause the specimens to be incorrectly assigned to a single species.

The K/θ method complements the ANI method of species delineation. ANI-based species cutoffs rely on ANI having a bimodal distribution with one cluster of interspecies ANI values, a second cluster of intraspecies ANI values, and a gap in between (e.g., Fig 3 in ref [28]). This empirical observation is predicted by the population genetic theory that underpins the K/θ method. Moreover, the K/θ method may be used to validate ANI cutoffs or as a second method for cases where ANI values overlap the cutoff (e.g., 95%) and thus do not support clear species separation.

### Multiple comparisons

We used a probability of 0.05 for hypothesis testing, understanding that when this is applied to many different cases, we expect to make a wrong decision in approximately 5% of the cases. This could be accounted for by using a correction such as the Bonferroni. On the other hand, many of the results are significant with a much higher probability. We choose not to use a correction and simply accept the risk of a small number of wrong conclusions, less than one in twenty for the cases in this paper.

### When small sample sizes can provide strong conclusions

Rosenberg [20] showed that small sample sizes can provide strong evidence that the entire populations from which they were taken are reciprocally monophyletic. This is because the probabilities are contingent upon the samples themselves being reciprocally monophyletic. Of course this assumes that the samples are representative of the populations, and do not, for example, come from a local sub-population that happens to be very different from the rest of the population. Thus one must always bear in mind the possibility that the specimens being studied are not representative. On the other hand, the possibility of erroneously splitting a species into two species based on a non-representative sample is likely to be small when K is very large. This is because, if a large K gives a K/θ value less than 4, the *θ* value must also be much larger than the generally observed values of < 0.05. There are parallels in traditional taxonomy, where new species are often described on the basis of one or a few specimens, especially when the new specimen is very different from the most similar described species. In this case we usually assume that additional new specimens will not have intermediate phenotypes that will cause us to combine the new specimen with a described species. If we had a single specimen of an African lion and two specimens of an ocelot, traditional taxonomy would probably recognize them as different species based on the absence of intermediate phenotypes, even without additional evidence. If gene sequences from these specimens had K/θ ≥ 4, this would indicate an absence of intermediate sequences and we would have similar objective evidence that they were different species.

### Use of trees and choice of tree-making methods

When implementing the K/θ method, the use of phylogenetic trees is limited to (i) finding sister pairs of clades for comparison; (ii) guiding the arrangement of sequences in the Excel spreadsheet to facilitate the calculation of K/θ; and (iii) spotting obvious cases of K/θ based on branch lengths. We use Neighbor-joining for tree construction because it is fast and easy to do with PAUP*, but presumably a different distance-based method could be used.

### Genes or phenotypes in bacterial species descriptions?

The bacterial rules of nomenclature do not specifically require phenotypic data. However, the report of an *ad hoc* committee on species definition in bacteria [29] calls for the use of qualitative phenotypic traits as well as 16S rDNA sequences to delimit species. A quick survey of some recent papers in the *International Journal of Systematic and Evolutionary Microbiology* shows that while rDNA sequences are now commonly used in descriptions of new bacterial species, they are used essentially like a collection of traits or to form phylogenetic trees. When these trees, or pairwise sequence differences, or both are used to define trees, there is usually no attempt to apply a theoretically justifiable method to distinguish sequence differences between species from differences within species. Instead, this is based on the judgment or intuition of the investigators.

### One gene or many?

There is a tendency in the recent literature to extoll the virtues of using many genes to discover species. But using more than one gene has several drawbacks. First, it costs more and will not be useful if one gene is sufficient to show that two samples come from different species. Second, if one gene is a core gene, adding another gene which is not a core gene and is involved in HGT can obscure the signal of speciation.

Ideally, to discover closely related species we would like to use the gene or genes that segregated earliest in those particular populations; in this way we would catch speciation in its early stages and discover closely related species. This is a real advantage to using multiple gene sequences, which would increase the chance that at least one of them would have segregated early. How important this is depends on the amount of sexual reproduction in the bacteria in question. At one extreme, if there is little or no recombination, or extreme inbreeding, all genes will segregate at the same time and any core gene should be as good as any other. At the other extreme, if there is extensive outcrossing and recombination, different genes may segregate at very different times after two populations begin evolving independently. In fact, a given gene may segregate early in one such population and late in the sister population. This is a likely explanation for the observation that CF1 and CF5 form a separate species from CF2, 3, and 4 for all protein-coding genes except *ompA*. Stackebrandt et al. [29] speak favorably of using protein-coding genes, claiming that there is a consensus that at least five such genes are necessary, but there is no such consensus in the sense of a number that is agreed on or at least accepted by nearly everyone in the field. More importantly, Stackebrandt et al. [29] do not consider the possibility that these genes may be at different stages of segregation, which is the simplest explanation of the different trees supported by different genes in the Wertz et al. [23] data.

There is also a potential problem with concatenated gene sequences: a significant K/θ ratio in one gene may be obscured by a small, insignificant K/θ ratio in another gene. In other words, the signal of species divergence in one gene that segregated early after speciation may be swamped by the signal in another gene that has not yet completed segregation with 95% confidence.

We suggest that the best method for large-scale diversity studies would be to use a single protein-coding gene to discover species and then, for groups of special economic, medical, ecological, or evolutionary importance or interest, test additional genes to see if they separate more species pairs because they segregated earlier in those pairs. Then it would often be useful to look for phenotypic or molecular markers that could be used to identify those species without requiring sequencing.

### Stability of EG species

A nice feature of species discovery using K/θ ≥ 4 is that additional specimens are unlikely to fill in the gap between species and require them to be lumped. That is because the large difference between K and θ reduces the probability that other specimens that are intermediate between the two species and fill in the gap between them will ever be discovered. Put another way, it is unlikely that we will find specimens and lineages that join the two clades we identify as species. If this did happen, the single species produced by joining the two clades would have exceptionally large nucleotide diversity.

Consequently, once two populations are assigned to different species using K/θ ≥ 4, those species are unlikely to be lumped in the future. It is perhaps more likely that data from additional genes or from new specimens will cause a species to be split in the future, and this has been seen repeatedly in the literature. But this is true for species delimited by any method, using any kind of data; in science, new information often forces us to change our conclusions.

### Other methods used to discover bacterial species with single loci

Most of the methods proposed for using DNA sequence data to delimit species are global methods, i.e. in a phylogenetic tree of gene sequences from many specimens, they look for the transition point from species divergence to divergence between individuals within species. One of these, GMYC [16] uses coalescent theory to detect the transition point. Barraclough et al. [30] used GMYC to discover species from environmental 16S rDNA sequences of bacteria but the method has not been applied to sequences from already-described taxonomic species of prokaryotes. Poisson Tree Picking (PTP) [31] attempts to identify significant changes in the numbers of substitutions along branches, which are expected to be significantly higher between species than within species. To our knowledge, PTP has not been applied to prokaryotes.

Ecosim [32] detects species produced by selective sweeps during adaptation to different habitats. It assumes a Poisson model of substitutions, in contrast to K/θ and GMYC that allow more complex and presumably more realistic models to be used. The theory behind Ecosim includes random drift and selective sweeps but not less drastic forms of selection. AdaptML [33] is another local method. It identifies clades associated with ecological parameters that might be relevant to ecological species. It detects species only if they are adapted to different niches that have already been identified. These methods will not detect species that evolve independently because they are physically isolated as in the case of *Sulfolobus islandicus* in hot springs [34] unless they are also adapted to different niches. In other words, they may fail to detect some evolutionarily independent species.

These global methods are all subject to error if different species have different levels of within-species variation, i.e. different values of nucleotide diversity. This is in contrast to K/θ ≥ 4 which is a local method focusing on sister clades and thus on putative sister species. Such pairwise comparisons have the advantage that sister species are more likely to have similar values of θ. In fact, the problem of delimiting species in general really is a local problem. We have no problem assigning specimens of the grizzly bear and the mule deer to different species, but it is more difficult to decide whether the Alaskan brown bear and polar bear are really different species, e.g. [35].

### Limitations of the K/θ method

In order to study the diversity of cyanobacteria from hot springs in Yellowstone National Park, Ward et al. [36] did environmental sequencing of clone libraries from the 16SrRNA/ITS region from PCR-amplified DNA. Their Figure 4 shows a neighbor-joining tree of ITS sequences of bacteria identified as *Synechococcus*. They found 10 putative ecotypes of *Synechococcus*, which are thus different species adapted to different niches. However, it is evident by inspection of the tree that only the three clades called A, A’, and B satisfy the K/θ ≥ 4 criterion. This could happen if most of the ecotypes have diverged too recently to pass the the K/θ ≥ 4 criterion for independent evolution. In this case ecotype analysis can divide species more finely because it delimits species at an earlier stage in the speciation process.

### Using K/θ in Formal Descriptions of New Species

We have shown that the K/θ ratio can be used to delimit evolutionary species in bacteria, illustrating the procedure with an exemplar data set. However, this does not mean that the species so delimited conform to current practice in bacterial systematics, or that the species so detected can be formally described in conformance with the current rules of taxonomic nomenclature for bacteria. Of course the same is true of species delimited with the GMYC method that also detects EG species, or with the method of Cohan [37], which detects a subset of EG species that are separated by ecology. Unfortunately, the rules of bacterial systematics codified in the International Code of Nomenclature of Bacteria are not explicitly based in population genetic theory, even though they pay lip service to the BSC which defines species as reproductively isolated populations in outcrossing sexual organisms.

### Response to the critique of Barraclough [19]

Barraclough [19] claimed to have found two errors in Birky’s [1-5] calculation that a K/θ value of 4 or greater is strong evidence of samples belonging to two different evolutionary species. First, citing Edwards and Beerli [38], he claimed that “K from the genealogy overestimates the true divergence time…due to sampling of the polymorphism from the ancestral common ancestor species” and that K is equal to 2Tμ plus the θ of the ancestor. However, Edwards and Beerli said that the addition of θ is necessary when K denotes the time elapsed since two populations diverged, but it is not appropriate when estimating the time T since allelic divergence, as we have done here and Birky did in previous papers using K/ θ. Second, Barraclough refers to “a mitochondrial marker with θ = Nμ .” In correspondence, he said that in this section he used N to refer to the effective population size rather than the census size, so he is claiming that θ = N_e_μ If this were true, then the upper 95% confidence limit of K/ θ would be 8. But by definition θ = 2N_e_μ [39]. This is because the sequence difference between two gene copies in two specimens A and B from the same species sharing a most recent common ancestor O is equal to the sequence difference N_e_μ between A and O plus the sequence difference between B and O, which is also N_e_μ. Consequently, after 4N_e_ generations, K/ θ = 8N_e_μ/2N_e_μ = 4, and not 8.

### Could we use K/ θ as a measure of the progress of speciation?

If we adopt the evolutionary species concept, then speciation is usually a gradual process. (Exceptions can include rare dispersal to isolated sites such as islands, or speciation by hybridization or polyploidization.) To the extent that speciation is gradual, the K/ θ ratio is expected to gradually increase. Consequently, if we have two populations that we suspect are undergoing speciation, K/ θ might be used as a measure of the progress of speciation. It would presumably be non-linear, as suggested by the nonlinearity of progress toward reciprocal monophyly of the two populations [20].

### Species are universal

In this paper and previous papers in this series [1-6] we have shown that Evolutionary Genetic species are found in bacteria and in both asexual and sexual eukaryotes. The arguments in this paper should also apply to Archaea. We strongly suspect that viruses also form such species, and that extraterrestrial organisms, if they exist, will likewise form populations that evolve independently from each other.

## Author contributions

CWB was responsible for study design, data collection and analysis, decision to publish, and preparation of the manuscript. HM assisted with manuscript writing and figure preparation.

## Acknowledgements

We thank John Wertz for providing the aligned sequences.

